# Genome-wide analysis of somatic non-coding mutation patterns and mitochondrial heteroplasmy in type B1 and B2 thymomas

**DOI:** 10.1101/2024.08.09.607250

**Authors:** Kohei Fujikura, Isabel Correa, Susanne Heck, Juliet King, Emma McLean, Andrea Bille, Daisuke Nonaka

## Abstract

**Introduction:** Type B1 and B2 thymomas are lymphocyte-rich malignant tumors with few somatic mutations in protein-coding regions of the nuclear genome; nonetheless, non-coding regions remain uncharacterized. Here, we developed a rigorous tumor isolation method from lymphocyte-rich thymoma tissues and identified somatic mutations in non-coding and mitochondrial DNA.

**Methods:** CD205^+^CD45^-^ pure tumor cells were isolated from fresh-frozen tissues using DEPArray system. Deep whole-genome sequencing was performed, and recurrent somatic alterations in coding, non-coding, and mitochondria regions were systemically identified by computational framework. The mutations were classified according to gene function, cis-regulatory element, and mutational signature.

**Results:** The total number of somatic mutations was approximately 80 times higher in non-coding regions than in coding regions in type B1-2 thymomas (1,671.3 vs. 21.1 per case). Coding mutations were identified in epigenetic regulators, DNA repair genes, and some other genes. Nevertheless, 40% of cases exhibited fewer than four mutations in coding regions. A systematic non-coding analysis identified a total of 405.0 mutations per case on cis-regulatory elements, and detected six recurrent mutations: one interferon regulatory factor (*IRF8*), two E3 ubiquitin ligases (*UBR2* and *RNF213*), and three intergenic regions. Mitochondrial heteroplasmy was observed in 90% of cases, with a significant proportion of mutations located in D-loop region. The single-base substitution pattern was signature 12.

**Conclusions:** Numerous non-coding mutations and mitochondrial heteroplasmy were detected in type B1 and B2 thymomas. Given the paucity of coding mutations observed in this disease entity, disruption of the non-coding landscape and mitochondrial heteroplasmic shift may be the primary cause of thymoma.

## Introduction

Thymomas are a type of thymic epithelial neoplasm that exhibit organotypical features observed in either the active or the senescent thymic gland.^1^ According to the World Health Organization (WHO) classification, they are divided into three types: A, B, and AB (WHO Classification of Tumours Editorial Board. Thoracic Tumours. Lyon (France): International Agency for Research on Cancer; 2021. (WHO classification of tumours, fifth ed.; vol. 5).^2, 3^ Type B is further subdivided into types B1, B2, and B3 based on the relative proportion of the non-tumorous lymphocytic component, and the resemblance to normal thymic architecture.^4, 5^ It has been demonstrated that different histologic subtypes of thymomas have distinct prognoses and clinical features.^6, 7^ For instance, type B thymomas have been found to have a higher stage (20-50% in stage III-IV) than type A and AB thymomas (90% in stages I–II).^6^ In addition, a greater incidence of autoimmune diseases is observed in patients with type B thymoma than in patients with type A thymoma.^7^ Surgery is the standard treatment for localized tumors, with postoperative radiotherapy and chemotherapy reserved for advanced stages.^8^ However, effective postoperative treatment has not been developed, possibly due to a lack of basic and clinical research compared to other malignancies.

Recent genomic studies of thymic epithelial tumors have revealed that type A and AB thymomas are characterized by a high frequency of thymoma-specific codon mutations (L424H) in the *GTF2I* gene.^9, 10^ On the other hand, no recurrent genetic abnormalities have yet been identified in the type B group, although a *KMT2A-MAML2* gene fusion has been reported in a small subset of type B2-3 thymomas.^11^

It has been suggested that B1 and B2 thymomas exhibit a markedly reduced number of somatic mutations in the protein-coding regions of the nuclear genome in comparison to other malignancies.^10, 12^ Thus, it is plausible that mutations in non-coding regions play a pivotal role in the pathogenesis of thymomas, as has been demonstrated in certain cancer types.^13–16^ Nevertheless, the non-coding regions of thymomas remain uncharacterized. In addition, type B1 and B2 thymomas contain a significant proportion (50-95%) of non-neoplastic immature T lymphocytes. The presence of a large non-neoplastic component renders genetic analysis of thymomas exceedingly challenging. In order to overcome these challenges and carry out genome-wide analysis, we developed a method for isolating pure tumor cells from frozen tissue sections. Subsequently, we conducted deep whole-genome sequencing and employed computational methods to identify significantly mutated non-coding regions and mitochondrial DNA.

## Materials and methods

### Ethics approval and consent

The study was approved by the Research Ethics Committee of King’s Health Partners Cancer Biobank (KHP Cancer Biobank REC reference 18/EE/0025). All procedures conducted in the study were in accordance with the 1964 Declaration of Helsinki and its later amendments or comparable ethical standards. Written informed consent was provided by all patients before the treatment procedure was commenced.

### Patients

Type B1 and B2 thymomas were retrieved from the King’s Health Partners Cancer Biobank. All tumors were treated by surgical resection for curative intent, without prior adjuvant therapy. The diagnosis was confirmed by reviewing Hematoxylin and Eosin (H&E) stained sections from resected tumors in accordance with the 2021 World Health Organization classification of thymic tumors.^2, 3^ The clinicopathologic information of the patients is presented in Supplemental Table 1. The median age of the patients at the time of surgery was 51 years (range, 29-79 years), with seven males and three females. Three patients exhibited type B1 thymomas, while seven exhibited type B2 thymomas. Six patients presented with paraneoplastic disease, including five with myasthenia gravis and one with autoimmune hepatitis. At the time of diagnosis, eight patients (80%) were staged as Masaoka stage I-II, while two (20%) were as for stage III-IV. The median tumor size was 60mm. Five patients (50%) had a positive margin (R1) and received postoperative radiotherapy for their primary tumors. One patient experienced recurrence. All patients demonstrated no evidence of other malignancies.

### Frozen tissue processing

Frozen sections (60 µm thickness) were fixed in 1 mL of 2 % paraformaldehyde for 15 minutes at room temperature. The sections were then washed four times with 1 mL of PBS and centrifuged at 800 x g for 5 minutes. The samples were digested in a solution of 5 mL of a mixture of collagenase and dispase in RPMI 1640 for 15 minutes at 37C, followed by incubation period of 15 minutes at 37C. Subsequently, 5 mL of 10 % FCS RPMI 1640 were added together with EDTA to a final concentration of 2 mM. Then, the samples were centrifuged at 460 x g for 10 minutes, after which the pellets were resuspended in 2 mL of FACS buffer and filtered through a 35 µm mesh. The samples were centrifuged at 460 x g for 10 minutes and resuspended in 1 mL of FACS buffer. The cell number was then counted, and a total of 1-2x10^5^ cells were stained in 100-200 μL of FACS buffer. After incubation with Fc block (Biolegend, CA) for 10 minutes at room temperature, APC anti-human CD45 (clone HI30; BioLegend) and PE anti-human CD205 (clone HD30; BioLegend) antibodies were added and incubated for 20 minutes at 4C. After 20 minutes of incubation, 2 μL of a 0.1 mg/mL stock of Hoechst (ThermoFisher, MA) were added to label the nuclei and samples were incubated for another 10 minutes at room temperature. Subsequently, the samples were washed with 2 mL of FACS buffer and pelleted at 460 x g for 10 minutes. The cell pellets were resuspended in 200 μL of FACS buffer.

### Cell isolation

The isolation of normal and tumor cells was conducted using a DEPArray V.II system with A300K cartridges (Menarini Silicon Biosystems, PA). A total of <1x10^5^ stained cells in FACS buffer were washed with 1 mL of DEPArray Buffer (Menarini Silicon Biosystems) and spun at 460 x g for 10 minutes. The cell pellet was washed once more with 1 mL of degassed buffers and was resuspended in 30 μL of the same buffer. The number of cells was counted and volume was adjusted to yield approximately 10,000 cells in 14 μL. The samples were then loaded into the cartridges according to the manufacturer’s instructions. The selected cells were eluted as single cells or in groups using 200 μL PCR tubes for collection. The PCR tubes were centrifuged at 14,100 x g for 30 seconds, after which 100 uL of PBS were added to the tube and centrifuged at 14,100 x g for 10 minutes. Subsequently, the supernatant was removed, leaving 1 μL of PBS with the pellet. Genomic DNA was amplified using the Ampli1 WGA kit (Menarini Silicon Biosystems) according to the manufacturer’s instructions. The quality of the amplified DNA was evaluated using the Ampli1 WGA QC kit (Menarini Silicon Biosystems). Only samples exhibiting four distinct bands were used for the genome sequencing. The final quantification of the purified samples was conducted using the Qubit fluorometer reagents (ThermoFisher) in the Glomax Discover plate reader (Promega, WI).

### Whole genome sequencing

The whole genome sequencing (WGS) library was constructed using the NEBNext Ultra™ II DNA Library (New England Biolabs, MA). The sequencing libraries were then multiplexed and clustered on the flowcell. Subsequently, the flowcell was loaded onto the Illumina NovaSeq 6000 instrument. The samples were subjected to sequencing using a 2x150 paired-end configuration. Image analysis and base calling were conducted by the NovaSeq Control Software on the NovaSeq instrument. The raw sequencing data (.bcl files) generated from the Illumina NovaSeq were converted into fastq files and de-multiplexed using Illumina’s bcl2fastq software. A total of at least 288 gigabases of raw read data was generated for all samples.

### Read mapping and detection of somatic mutations

The raw sequencing data were processed using the DRAGEN DNA Pipeline on the Illumina DRAGEN Bio-IT Platform v4.1.7 (Illumina). Following adapter trimming, the sequencing reads were aligned to GRCh38/hg38. Subsequently, mutations were filtered based on the following criteria: (1) "PASS" in the quality filter; (2) Somatic Qscore ≥18; (3) tumor sample coverage ≥ 21X; (4) normal sample coverage ≥ 21X; (5) tumor frequency > 0.25; (6) ≥ 6 distinct reads supporting the mutation in the tumor sample; (7) normal frequency < 0.02. Both germline and somatic exomes were analyzed simultaneously, and variants present only in somatic DNA were extracted using the variant caller. The variants in exonic regions were classified into six categories: synonymous, non-synonymous, splice acceptor, splice donor, stop-causing, and stop-loss variants. Some sequencing reads in coding regions containing variants were visually inspected using the Integrative Genomics Viewer (IGV) (http://software.broadinstitute.org/software/igv/) to exclude potentially artifactual variants as described previously.^17, 18^ These included variants occurring in variant-rich regions or variants identified exclusively at read ends.

A list of previously reported non-coding mutations in different cancer types was retrieved from ten international non-coding projects, which clearly define the chromosomal positions. The following table lists the cancer types and original data references:

> Pancancer projects, reference: [13] [14] [19] [20] [21] [22]^13, 14, 19–22^

> Hepatocellular carcinoma project, reference: [15]^15^

> Breast cancer project, reference: [16]^16^

> Urinary bladder cancer project, reference: [23]^23^

> Diffuse large B-cell lymphoma project, reference: [24]^24^

De novo detection of recurrent mutation was performed using the MutSpot R package^25^ with the default settings. Manhattan plot was generated based on the chromosomal position and *P*-value.

### Tumor mutational burden

Tumor mutational burden (TMB) was defined as the number of somatic, coding, base substitution, and indel mutations per megabase of genome examined. The calculation encompasses all base substitutions and insertions (indels) in the coding region, including synonymous alterations. Non-coding alterations were not included in the analysis.

### Characterization of coding mutations

The functional impact of an amino acid substitution on the protein structure and overall function was predicted using SIFT and PolyPhen. The gene ontology (GO) terms of molecular functions, cellular component, and biological processes of the detected mutations were searched using the web-based tool Metascape (https://metascape.org/gp/index.html#/main/step1). The mutated genes were analyzed in the STRING database (http://string-db.org) to evaluate their relationships through protein– protein interaction (PPI) information. The PPI information was imported into the Cytoscape software, which calculated the relationships between mutant proteins in type B1-2 thymoma. Statistical significance was determined by *P* < 0.05 and a false discovery rate (FDR) < 0.05.

### Characterization of non-coding mutations

The ENCODE Registry of candidate cis-Regulatory Elements (cCREs) DNA regulatory elements was used to identify DNA regulatory elements in the human genome. The ReMap 2022 Atlas of regulatory regions, which is an integrative analysis of all public ChIP-seq data for transcriptional regulators from GEO, ArrayExpress, and ENCODE was retrieved from the UCSC genome browser or ReMap2022 database (https://remap.univ-amu.fr/). The dplyr R package was used to ascertain whether the identified gene mutations were located on the cis-regulatory elements.

### Detection of mtDNA mutations

The raw sequencing data were processed using the DRAGEN DNA Pipeline on the Illumina DRAGEN Bio-IT Platform v4.1.7 (Illumina). After adapter trimming, the sequencing reads were aligned to GRCh38/hg38. Subsequently, a series of filter conditions were used to call mtDNA variant sites, including mutations were filtered based on the following criteria: (1) "PASS" in the quality filter; (2) Somatic Qscore ≥18; (3) tumor frequency > 0.05; (4) ≥ 5 distinct reads supporting the mutation in the tumor sample; (5) normal frequency < 0.05. Both germline and somatic exomes were analyzed simultaneously, and variants present only in somatic DNA were extracted using the variant caller. The heteroplasmy level was defined as the proportion of mutant reads in the total reads for a given mutation site. Our analysis demonstrated that sequencing coverage had no notable effect on heteroplasmy level, thereby providing compelling evidence of the high technical reliability of the method.

### Detection of copy number alterations

The CNV workflow provided with the DRAGEN Somatic Platform (Illumina) was performed based on 150 bp fragments. The reads were counted in 500 kb bins, followed by GC bias correction and normalization based on a reference set. Segmentation was conducted via Shifting Level Models with the disabled merging of two adjacent segments. The interactive Circos plot was generated using the R-package shinyCircos.

### Analysis of scRNA-seq thymoma data

The publicly available data for the scRNA-seq dataset of thymoma samples were obtained from the SingleCellPortal (https://singlecell.broadinstitute.org/single_cell). The thymoma study included four primary tumor and blood samples. The blood samples and type AB thymoma samples were excluded from our study. The expression levels (transcripts per million, or TPM) were obtained, with tumor cells identified using previously determined criteria. Uniform manifold approximation and projection (UMAP) representations of the cells were generated from the total expression profiles. The expression (TPM) values were then compared between the given factors.

### Immunohistochemistry

Deparaffinized sections were heat-treated and incubated with the primary CD205 antibody (clone: 11A10, dilution 1:25; Novocastra, Newcastle, United Kingdom). Immunostaining for CD205 was performed on a Bond Max autostainer (Leica Microsystems, Wetzlar, Germany) according to the manufacturers’ instructions.

### Statistical analysis

All statistical tests were carried out using GraphPad Prism version 10 for Windows (GraphPad Software, San Diego, CA). The Mann–Whitney U test or the Wilcoxon signed-rank paired test was used to compare the difference between two groups with continuous variables. The Fisher’s exact test was used for discrete variables. All P-values were calculated using two-tailed test, and statistical significance was set at the level of *P* < 0.05.

## Results

### A strategy to analyze type B1 and B2 thymoma genome

To comprehensively detect genome-wide mutations in non-neoplastic lymphocyte-rich tissues, we aimed to develop a rigorous tumor isolation method that preserves genomic quality and reduces various biases. Firstly, we analyzed publicly available data on scRNA-seq datasets of thymoma samples to determine the specificity of existing tumor-specific markers in type B1 and B2 thymomas. The scRNA-seq data demonstrated that thymoma tissue is composed of large T cell, B cell, and myeloid cell clusters, and a very small number of stromal cells, including tumor cells (Figure 1A). CD205 (DEC205/LY75) is a specific surface marker for thymoma that we have proposed (Supplemental Figure 1A),^26^ and when this marker is combined with CD45, it is possible to strictly isolate minute amounts of tumor cells (Figure 1B and 1C and Supplemental Figure 1B). The theoretical percentage of CD45^-^CD205^+^ tumor cells was 0.14% for B1 thymoma and 0.36% for B2 thymoma in scRNA-seq dataset. The CD45^+^CD205^-^ cell population (>95%) was considered non-tumorigenic and best suited as a matched control. The CD45^+^CD205^+^ cell population was small and mostly dendritic cells. These results raise the question of whether some existing genetic analyses have been performed with sufficient tumor cellularity, and demonstrate the importance of our research strategy. Here we used the DEPArray image sorter to isolate the pure tumor cells (Figure 1D and 1E), and extracted genomic DNA for genome-wide identification of somatic alterations (Figure 1F).

**Figure 1.**
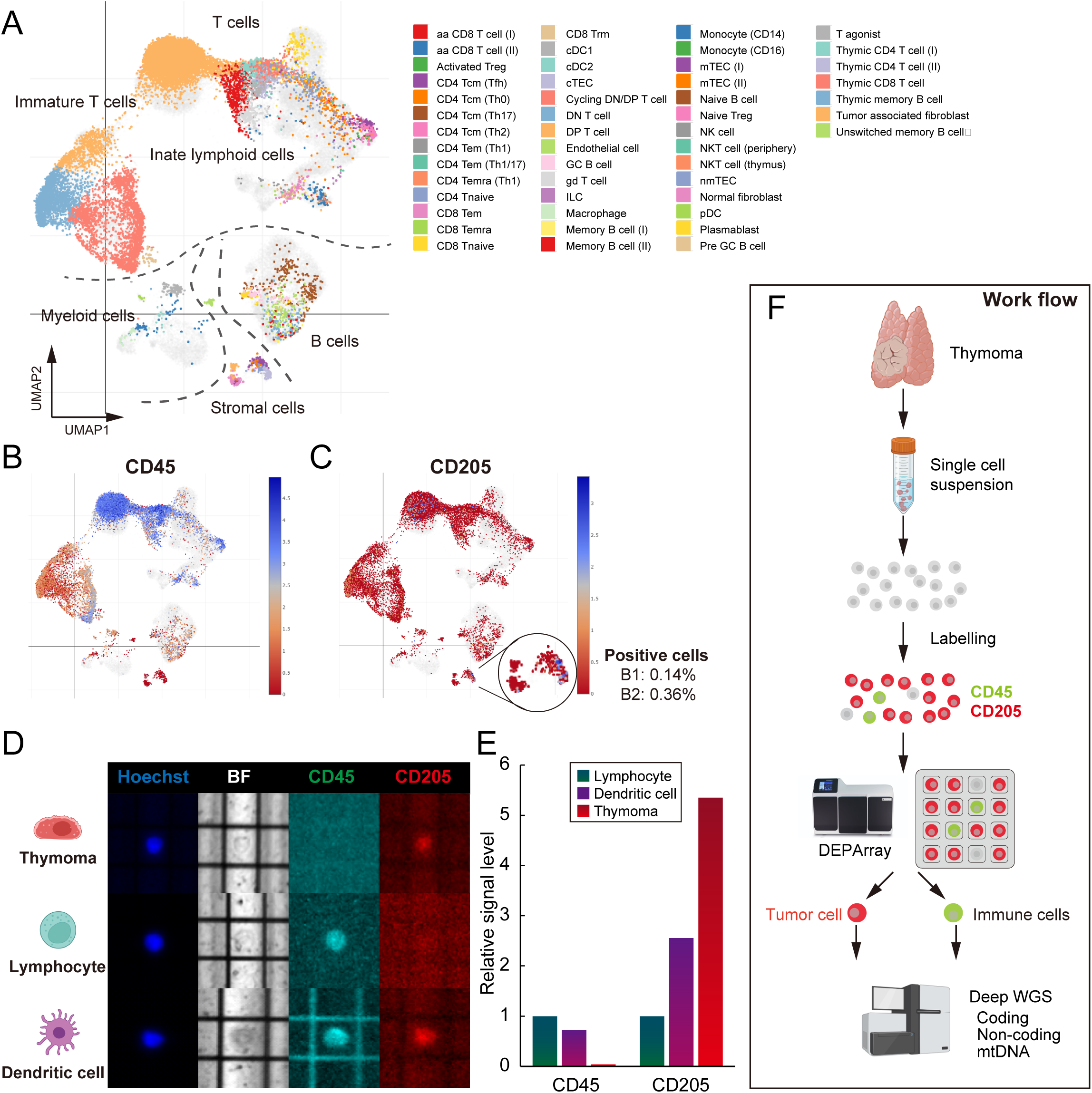
A strategy to isolate pure tumor cell. (A) Uniform manifold approximation and production (UMAP) plot displaying 49 clusters from thymoma patients (total 15,731 cells). (B, C) UMAP plot of marker CD45 (PTPRC) and CD205 (LY75). (D) Representative images of cells on the DEPArray. Cells were stained with CD45, CD205, and Hoechst for identification and selection. (E) Quantification of signal levels of CD45 and CD205 in each cell type. (F) The workflow to rigorously isolate thymoma and matched control cells for deep WGS.

### Characterization of type B1-2 thymoma genomes

Deep sequencing was performed across entire genome regions. Sequencing depths for tumors and corresponding normal components were 115× and 129× on averages, respectively, which were greater than those in typical WGS studies (Supplemental Table 2). Comparison of tumor cells with normal cells showed that all thymoma samples, including the pT3 samples, had a low TMB (mean 0.42 mutations/Mb in protein-coding region, range 0.03–0.97). The TMB of type B1-2 thymoma was the lowest among the various tumors, with 40% of cases having four or fewer mutations observed in the coding region (Figure 2A and 2B).

**Figure 2.**
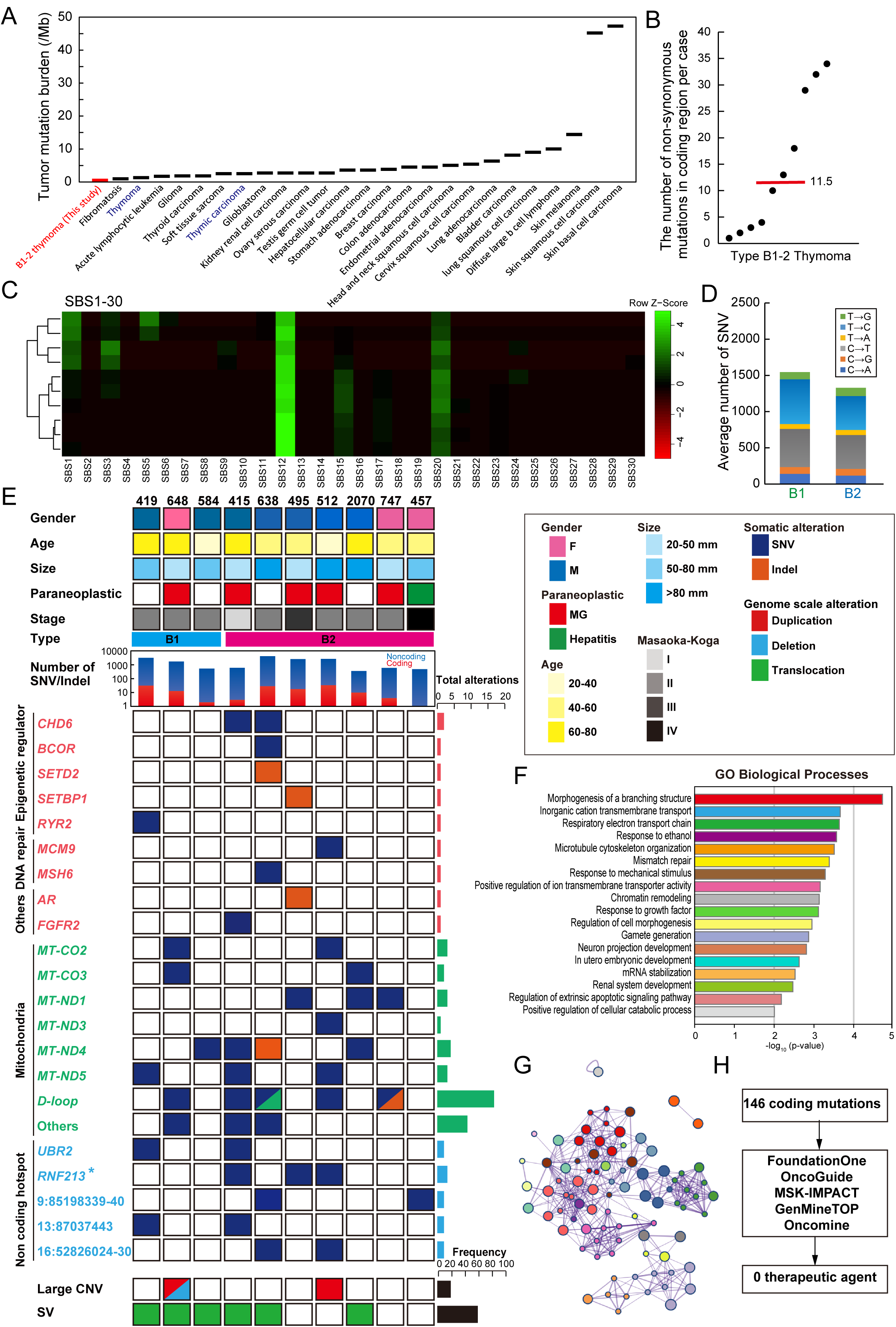
Identification of coding, non-coding, and mitochondrial mutations. (A) The frequencies of somatic mutations observed in protein-coding regions. The y-axis represents the median number of somatic mutations observed in each tumor, while the x-axis indicates the tumor type. Tumor types are ordered by the number of somatic mutations. (B) The distribution of coding mutation burdens per patient in type B1 and B2 patients. The number of somatic DNA variants per patient is depicted along the y-axis, with each dot representing an individual thymoma patient. The red-colored bar represents the median value. (C) Mutational signatures. Unsupervised clustered heatmap of contributions from 30 mutational signatures (SBS1-30) for thymomas. (D) Comparison of six base-substitution patterns between B1 and B2 thymomas. (E) Detection of somatic mutations in type B1 and B2 thymomas. The main plot shows information for coding and non-coding regions with mutations in 10 thymomas. Mutations most likely to be related to thymoma pathogenesis are classified into seven categories indicated at the left: I, epigenetic modifiers; II, DNA repair pathway; III, other genes; IV, Mitochondria genes; V, non-coding hotspot; VI, large copy number variations; VII, structural variants. Deep blue, SNV; orange, Indel; red, duplication; light blue, deletion; green, translocation. The upper histogram displays clinical information (i.e., gender, paraneoplastic disease, age, tumor size, Masaoka-Koga classification, thymoma type) and total number of SNVs and indels. Asterisk (*): Case #512 has a coding mutation while cases #415 and 495 have non-coding hotspot mutations. (F) Functional validation of frequently mutated genes in thymoma. The GO Biological processes were discovered by enrichment analysis for thymoma-associated genes. (G) Network analysis. The cancer-related genes found in our datasets were examined with the help of Cytoscape to identify the human-curated pathway datasets. The size of the spheres indicates the quantity of genes detected in our cohort, and the color of the spheres corresponds to the GO biological processes shown above. (H) Strategy to identify the potential therapeutic agents based on the current companion diagnostics.

The nucleotide substitution pattern of mutations assists in clarifying the background of tumorigenesis. Mutation signatures were successfully determined in all cases. The spectrum of base substitutions showed a strong predilection for T>C and C>T transitions, and it was similar to the known cancer mutational signature (single-base substitution 12: SBS12) reported by global cancer studies (Figure 2C).^27^ This pattern is reported in a small percentage (<20%) of hepatocellular carcinomas, but exact etiology is still unclear.

No specific differences were detected between type B1 and B2 thymomas with respect to the proportion of coding/non-coding SNVs, damaging/benign mutations, and mutation signatures (Figure 2D and Supplemental Figure 2).

### Paucity of coding mutations in nuclear genome

Exon analysis detected somatic mutations in epigenetic regulators (*CHD6*, *BCOR*, *SETD2*, *SETBP1*, *RYR2*), DNA mismatch repair genes (*MCM9*, *MSH6*), collagen-related genes (*COL4A1*, *COL6A3*, *COL6A5*), and some other genes (*AR*, *FGFR2*, etc) (Figure 2E and Table 3). GO analysis showed that the mutated genes were collectively involved in chromatin remodeling, DNA repair, stress response, organ development, and cell morphology (Figure 2F-G and Supplemental Figure 3). Gene mutations previously identified in type A and AB thymoma (e.g., *GTF2I*, *HRAS*, *KRAS*)^9, 10, 28–30^ were not found in our cohort. *TP53*, *KIT*, and *CDKN2A* are frequently mutated in thymic carcinoma,^9, 10^ but were wild type in type B1 and B2 thymomas examined. Using the gene companion diagnostic approach currently used in clinical practice, we attempted to identify potential therapeutic targets based on detected protein mutations. However, after applying five different panel testing algorithms, no candidate therapeutic agents were identified as of May 2024 (Figure 2H).

### Comprehensive profiling of non-coding mutations

The median number of exonic non-synonymous alterations per case was 11.5. 40% of cases exhibited a total number of four or fewer genetic mutations in the protein-coding region (Figure 2B). These findings underscore the necessity to focus on non-coding regions. To investigate the non-coding landscape of type B1-2 thymomas, we conducted a genome-wide survey of non-coding regions with an average coverage of 39.4× (range: 30.3×–65.1×) (Supplemental Table 2). The mean total number of somatic mutations in non-coding regions per case was 1,671.3 (range: 348-4,164), which were 79.2 (range: 29.0-484.0) times more frequent than in coding regions in type B1-2 thymomas (Figure 3A). The majority of non-coding mutations were observed in introns (41.6%) or intergenic regions (44.6%), yet the mutation rate per Mb were comparable across the six different functional regions (range: 0.45-0.64/Mbp) (Figure 3B).

**Figure 3.**
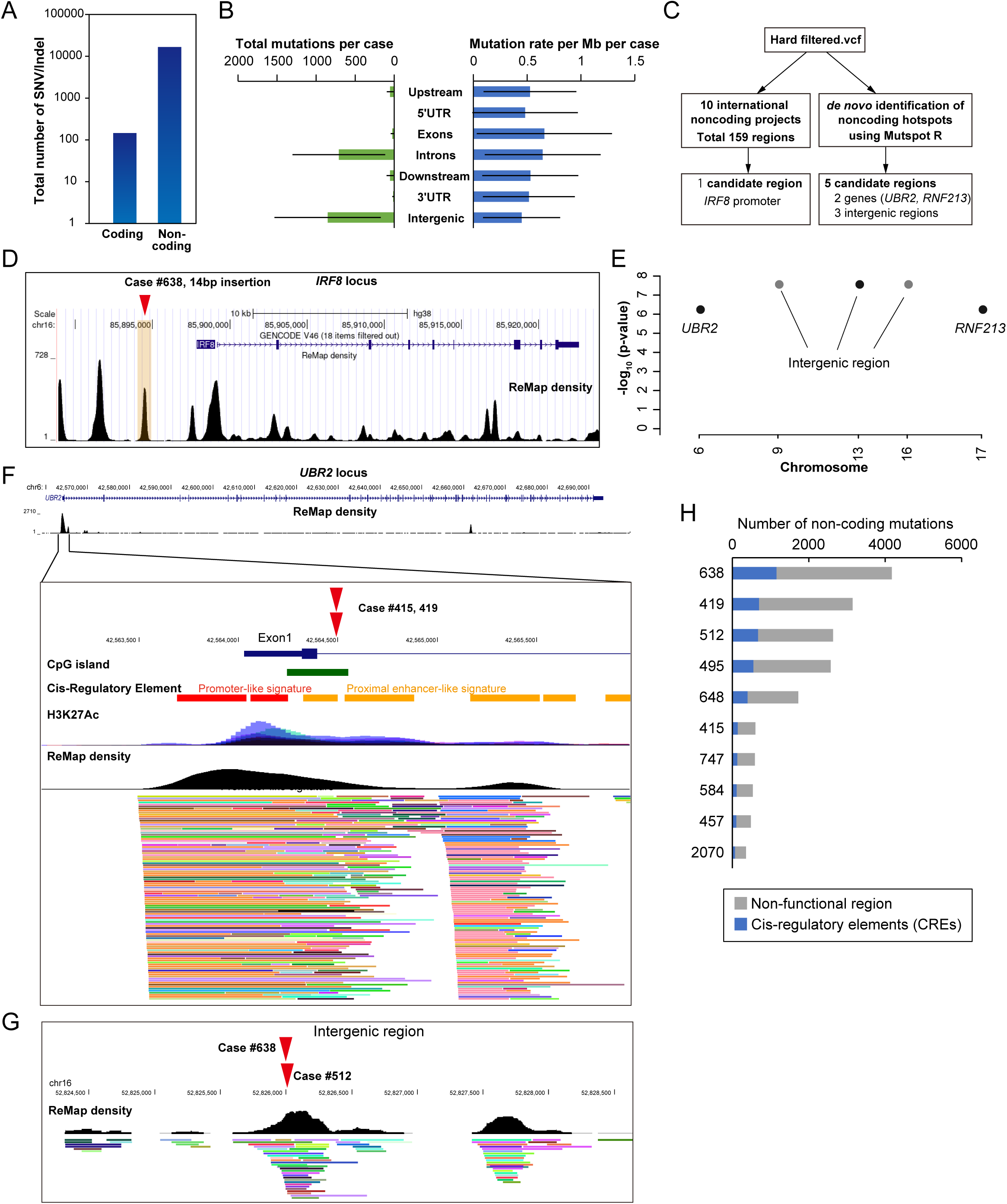
Identification of non-coding mutations. (A) Comparison of the total number of mutations in the coding vs non-coding regions. (B) Total number of mutation events (SNV + indel) and mutation rates (total events per Mb per patient) in different functional regions. (C) Strategy to identify the significantly mutated non-coding regions. (D) Identification of mutations in the *IRF8* promoter based on a list of known hotspot regions, which are detected in ten international projects. (E) Manhattan plot of the recurrent non-coding mutations. Figure is produced by MutSpot from a genome-wide analysis of whole genomes. Hotspots with FDR <0.05 are plotted. The mutations are located in the region characterized by Remap density (Black). (F) Non-coding mutations in *UBR2* locus and its genomic locus characterization. *UBR2* mutation is located in the enhancer characterized by CpG island and H3K27ac (Blue: gene structure, Green: CpG island, Red: putative promoter, Orange: putative enhancer, Purple and blue: H3K27Ac, Black: Remap density). (G) Mutations in intergenic region and its genomic locus characterization. The mutations are located in the region characterized by Remap density (Black). (H) The number of mutations in cis-regulatory elements. The non-coding mutations are divided into two regions: Non-functional region (gray) and cis-regulatory elements (CREs; blue).

Two distinct methodologies were employed to identify the recurrent non-coding mutations. In the initial approach, we conducted a search for the previously identified recurrently mutated regions, retrieved from ten international projects (Figure 3C). A total of 159 regions have been listed as significantly mutated regions, and *IRF8* promoter region was mutated in our study (Figure 3D and Supplemental Table 4).

In the second approach, we tried *de novo* identification of non-coding hotspots using MutSpot R-package on the entire WGS dataset (Figure 3C). This computational workflow identified five recurrent mutations: two E3 ubiquitin-protein ligase (*UBR2* and *RNF213*) loci and three intergenic regions (Figure 3E, Supplemental Figure 4 and Table 4). The mutation in the *UBR2* locus was identified in the CpG-rich enhancer in Intron 1 (Figure 3F). The mutations in the *RNF213* locus were identified in the last intron (Intron 67) in two cases and in the coding mutation (Exon 67) in another case (Supplemental Figure 4). The intergenic region on chr16 52826024-52826030 displayed regulatory signals, suggesting the potential significance of this region at the transcriptional regulatory level (Figure 3G). A systematic non-coding analysis identified a total of 405.0 mutations per case on putative cis-regulatory elements, while 1266.3 mutations were classified as non-functional (Figure 3H).

### Copy number and structural variants

Arm-level somatic copy number alterations were detected in 20% of cases (Figure 4, Supplemental Figure 5 and Table 5). An average of 2.3 fusion events were identified per patient, with 60% of patients having translocations, including 1.7 interchromosomal fusions and 0.6 intrachromosomal fusions (Figure 4, Supplemental Figure 5 and Table 6). None of these translocations have been previously reported. The *MAML2* gene fusions (*KMT2A-MAML2*, *YAP1-MAML2*, and *CRTC1-MAML2*), which have been observed in a small subset of thymic epithelial tumors, were not identified in our cohort.^11, 31–33^

**Figure 4.**
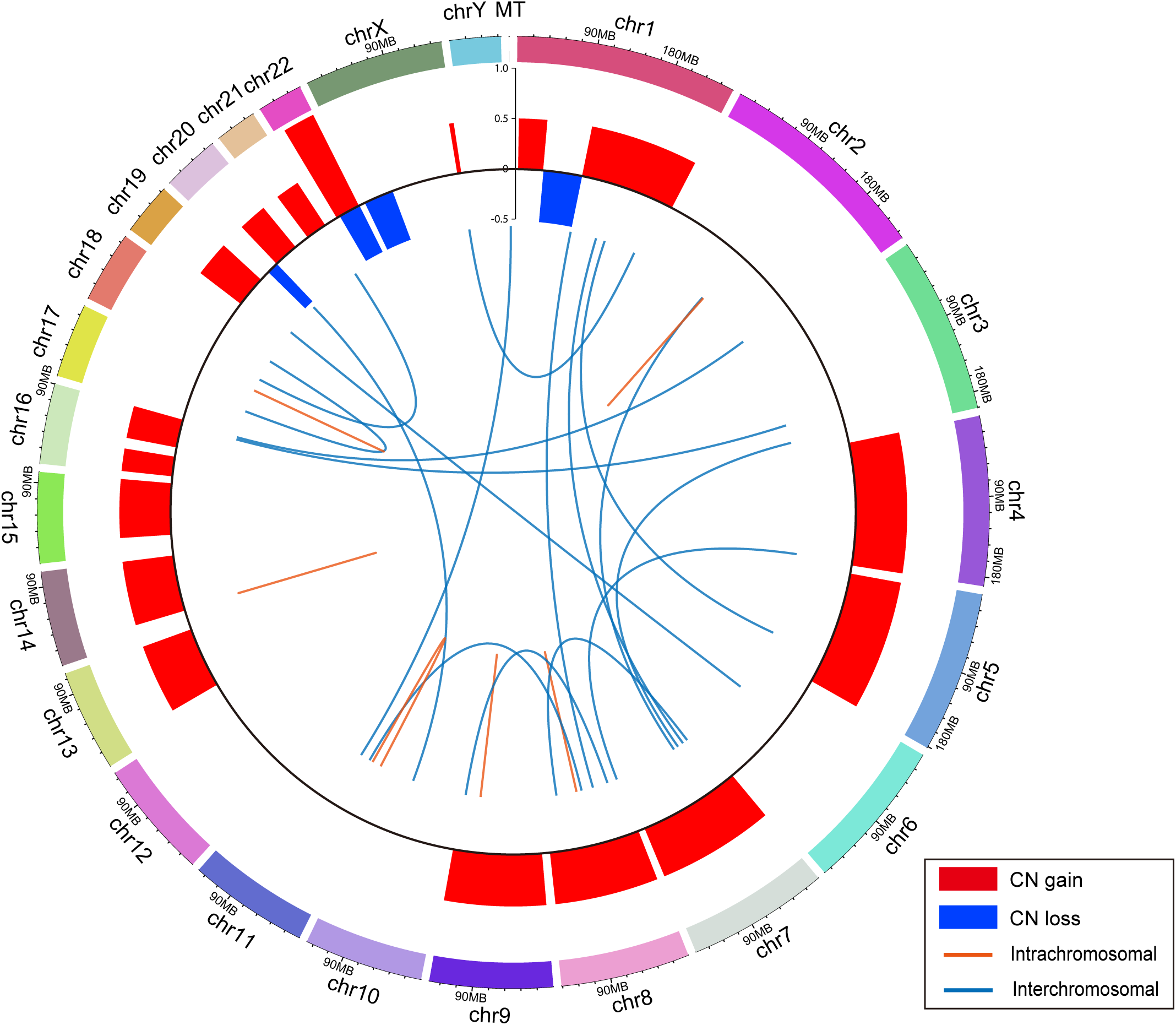
Genomics landscape of copy number and structural variants in type B1 and B2 thymoma. Circos plot for large-scale genomic alterations in 10 thymomas. Copy number events are summarized in the outer circle with red and blue color indicating copy number gains and losses, respectively. Translocations are marked by purple (interchromosomal) and green (intrachromosomal) lines; for intrachromosomal translocations, the green connecting line may appear as a single line if the joined regions lie within 1 Mbp.

### Mitochondrial heteroplasmic shift

In thymoma, somatic alterations in mitochondrial DNA (mtDNA) have not previously been focused on. We conducted a survey of mtDNA sequences with an average coverage of 368× (range: 71×–1244×) (Supplemental Table 2). An average of 4.3 mtDNA mutations were identified per patient, with 90% of patients having mutations (Figure 5 and Supplemental Table 7). The mutation density (number of mutations per length of kb) in mtDNA was approximately 500 times higher than that in nuclear genome (2.6/kbp vs 0.005/kbp) (Figure 5A). The mutation signature revealed a strong bias towards T>C transitions in mitochondria as well as in nucleus (i.e., SBS12 pattern), suggesting that similar mechanisms contribute to mutation accumulation between the nuclear and mitochondrial genomes (Figure 5B).

**Figure 5.**
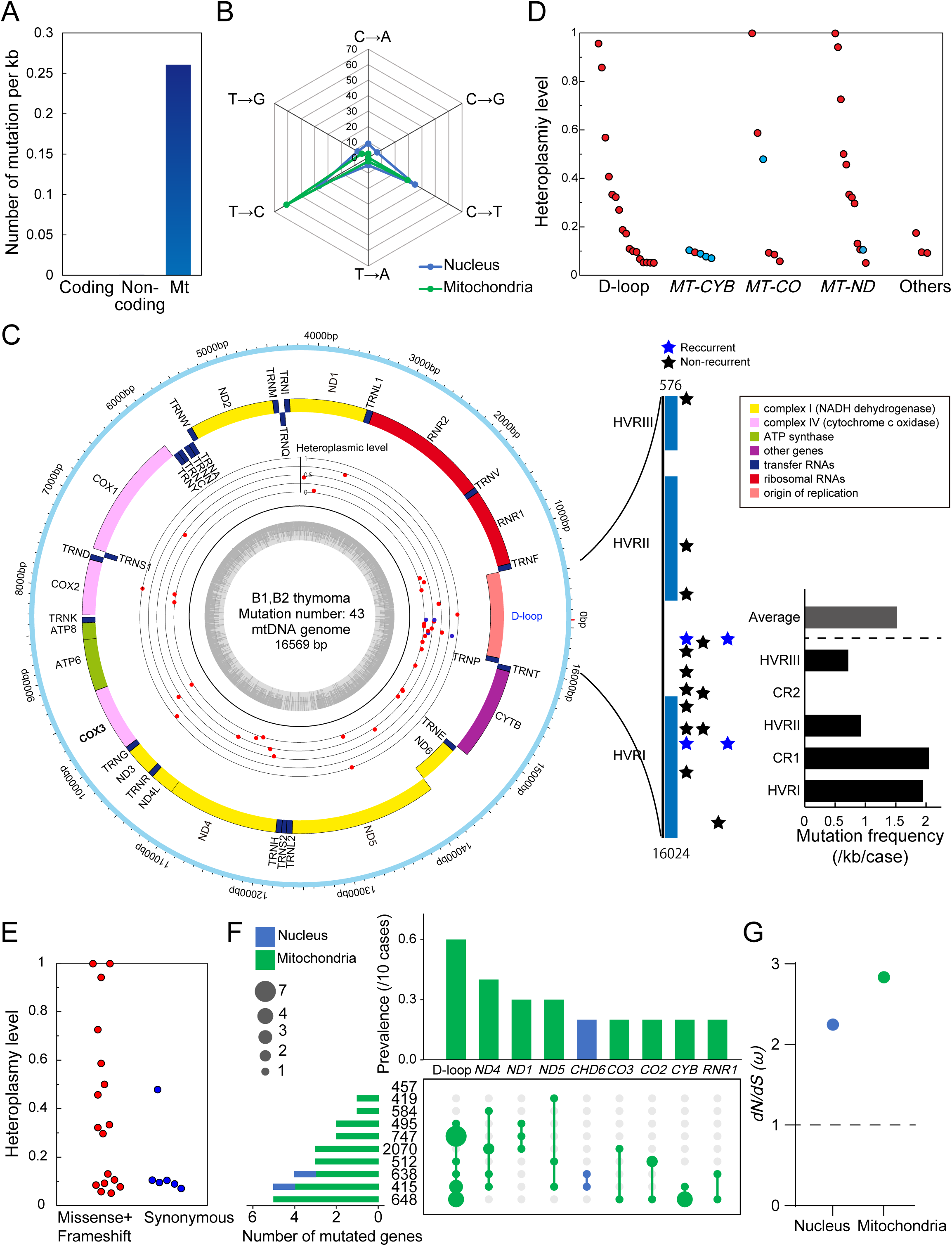
Somatic mtDNA mutations in thymoma patients. (A) Comparison of the somatic mutation rate in the coding vs non-coding vs mitochondrial regions. (B) Comparison of six base-substitution patterns between nuclear and mitochondrial genome. (C) Circos plots for the distribution of mtDNA somatic mutations in thymoma. Somatic mutation events are summarized in the inner circle with blue color indicating the recurrent mutations. Right vertical bar shows the distribution of D-loop mutations. HVR: Hypervariable region, CR: conserved region. (D) The heteroplasmy level of each mtDNA mutation in five different functional regions. (E) Comparison of the heteroplasmy levels of each mtDNA mutation between nonsynonymous and synonymous mutations. (F) A summary of genes commonly mutated in multiple cases. Upset plot of shows the prevalence of each gene mutation. Blue indicates the mutations in nuclear genome, while green indicates the mutations in mtDNA. (G) Comparison of *dN/dS* (ω) between nucleus and mitochondrial coding mutations.

**Figure 6.**
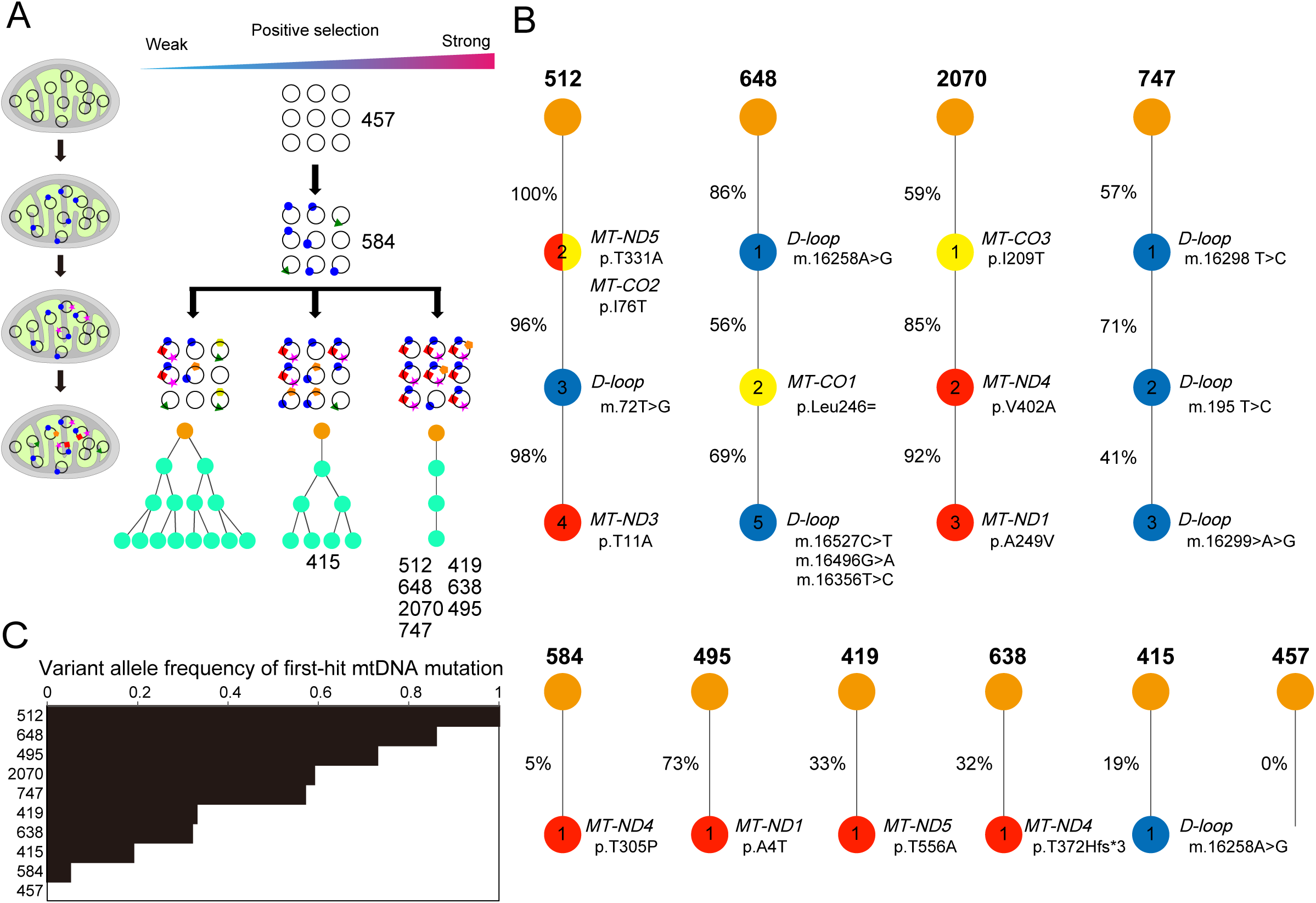

The D-loop region was most frequently mutated (n=17), followed by *MT-CYB* (n=5), *-ND4* (n=5), *-ND1* (n=3), *-ND5* (n=3), *-CO2* (n=3), and other regions (n=7) (Figure 5C and Supplemental Table 7). Hypervariable region I (HVR I) and its surrounding regions showed a high density of genetic mutations, with two recurrent mutations detected (m.16258 A>G and m.16527 C>T). The transmission of mitochondrial DNA to the nuclear genome (Chr 11) was also detected in tumor cells (Figure 4). Heteroplasmy levels were similar between different genes, but different between non-synonymous and synonymous mutations. A summary of genes commonly mutated in multiple cases revealed that only the *CHD6* gene was mutated in the nuclear genome, while eight genes were mutated in the mitochondrial genome (Figure 5F). The *dN/dS* ratio (ω) of protein-coding genes was greater than 1 in both mitochondria and nuclei, and was higher in mitochondria than in nuclei (Figure 5G).

## DISCUSSION

The comprehensive identification of somatic mutations that contribute to the tumorigenesis is a priority of tumor sequencing projects. To date, the majority of studies have focused on mutations within protein-coding regions of the genome.^34^ Nevertheless, several recent discoveries, including the identification of recurrent somatic mutations in the *TERT* promoter in various cancer types, indicate that mutations in non-coding regions are also crucial in tumor development.^35^ Moreover, the analysis of whole-genome sequencing data from tumors has elucidated novel mutational patterns and processes imprinted on cancer genomes.^27^

Thymoma is unique in that the number of coding mutations is very low,^12^ therefore it is essential to prioritize the analysis of non-coding regions in genomic research. Especially, type B1 and B2 thymomas are predicted to have low numbers of coding mutations.^10^ The genetic landscape of type B1 and B2 thymomas is less characterized compared to other cancers and other types of thymomas because genetic investigation is hampered by the abundance of non-tumorous immature T lymphocytes. Also, deep WGS analysis is five times more expensive than whole-exome analysis.^36^

In this study, we have developed the new methods that utilize an automated image cell sorter in combination with CD45 and CD205. Our method achieved 100% purity of tumor cells and the complete elimination of immature lymphocytes. Consequently, we believe that this methodology enabled molecular profiling of pure tumor cells from Type B thymoma tissues and can serve as a basis for thymoma research. The current limitations of this approach include the fact that the DEPArray system is not readily available at all institutions and that the experimental setup is not straightforward, time-consuming and labor-intensive.

We have newly identified five non-coding regions that are repeatedly mutated in type B1-2 thymomas. One of the identified regions is the *RNF213* locus, a large E3 ubiquitin ligase with a dynein-like core involved in a distinct ubiquitin transfer mechanism. The *RNF213* gene is mutated in some malignant tumors, including gastric cancer, ovarian cancer, lung cancer, liver and bile duct cancer,^37–39^ yet there have been few functional studies on *RNF213* mutations in malignancies. Previous reports have indicated that *RNF213* acts as a tumor suppressor and, its mutation promotes tumor development and progression.^37–40^

Another mutated region is the *UBR2* locus, a RING E3 ligase that regulates gene expression and chromatin-associated ubiquitylation in response to DNA damage. *UBR2* is widely expressed in a variety of human cancer tissues, particularly breast cancer, prostate cancer, and lymphomas.^41^ This gene has been demonstrated to promote tumor growth and metastasis.^41^ In addition to proteasomal degradation, *UBR2* has been shown to regulate the Erk/MAPK pathway, thereby preventing caspase-independent cell death.^42^ In this study, two E3 ubiquitin ligases were identified. Malignant tumor cells exhibit a high dependence on the proteasomes, and it may be possible to exploit this dependence to induce cell death using proteasome inhibitors.

It is widely recognized that mtDNA mutations play an important role in the initiation and progression of malignant tumors.^43^ However, there are no reports on mitochondrial heteroplasmy in thymomas and thymic carcinomas. This study provides the first systematic and comprehensive profiling of mtDNA mutations in thymoma cells, with the following advantages: (i) The average NGS coverage depth of more than 360-fold enables the accurate detection of the mtDNA mutations with heteroplasmy level as low as 5%; (ii) The data from matched control and thymoma tissues allows accurate identification of thymoma-associated mtDNA mutations; (iii) The DEPArray-based tumor cell isolation ensures the validity of mtDNA mutation profiling in thymoma tissues.

Despite the small size of mitochondrial genome, 43 mutations were detected. A significant proportion of these mutations were located in the D-loop, with heteroplasmy levels ranging from 5% to 96%. Consequently, our findings strongly underscore the pivotal role of the D-loop region mutations in the pathogenesis of thymic epithelial tumors. The significance of the D-loop has also been reported for several other malignancies, including breast cancer and hepatocellular carcinoma.^44, 45^ Previous reports have also indicated that D-loop mutations with high levels of intratumoral recurrence in cancerous tissues also have high levels of heteroplasmy. These findings indicate the possibility of positive selection for D-loop mutations in tumorigenesis. Similarly, high levels of heteroplasmy for mutations in the mtDNA coding region in thymomas also suggests the presence of positive selection. This is consistent with previous findings that cancer mutations predicted to have large functional effect are significantly enriched and can be distinguished from passenger mutations with low functional effect.^46^ This is also consistent with our result that the heteroplasmy levels are lower for synonymous than non-synonymous mutations. In conclusion, these results provide compelling evidence supporting the crucial role of mtDNA mutations, particularly those in the D-loop region, in thymoma tumorigenesis.

It has been reported that somatic D-loop mutations are significantly associated with reduced mtDNA content in malignant tumors and poor patient prognosis. Mouse with mitochondrial mutator phenotype, generated by a proofreading-deficient version of DNA polymerase gamma (*Polg*), exhibited an accumulation of mtDNA mutations, including D-loop mutations, and a reduction in mtDNA copy number.^47^ In addition, mutant *Polg* was shown to markedly enhance the invasive potential of breast cancer cells *in vitro*.^48^ Together with these results, the findings presented in Figure 5 indicate that the D-loop mutations may contribute to the malignant transformation of thymic epithelial cells by reducing mtDNA copy number. This hypothesis must be definitively confirmed by future development of mtDNA editing technology that can generate site-specific mtDNA mutations,^49^ particularly in the D-loop region.

The mutation signature was also investigated in this study. SBS12 was detected in all cases of type B1 and B2 thymomas. SBS12 has previously been reported in hepatocellular carcinomas and renal carcinomas.^27, 50^ A recent report has indicated that the proportion of SBS12 may vary from country to country.^50^ These findings indicate that exposure to agents that contribute to SBS12 mutations in liver and kidney cancer is prevalent in certain countries. The specific agent responsible for SBS12 remains unknown and requires further investigation. Nevertheless, the precedent set provided by other mutation signatures involving transcriptional strand bias suggests that the agent is probably exogenous.

In conclusion, numerous non-coding and mtDNA mutations were identified in type B1 and B2 thymomas. Given the paucity of coding mutations observed in this disease entity, it is possible that the disruption of the non-coding landscape may be the primary cause of thymoma. Targeting dysregulated signaling pathways and cellular processes involved in thymoma progression, such as ubiquitination and mitochondrial heteroplasmy, may be a promising approach for precision medicine in thymoma therapy. Furthermore, the combination of CD205 and CD45 with the DEPArray system was proved to be useful for genetic analysis of thymoma. Continued efforts to unravel the genetics of thymoma through our approach will advance our understanding of thymoma biology and improve patient care.

## Supplemental Figures and Tables

**Supplemental Figure 1.**
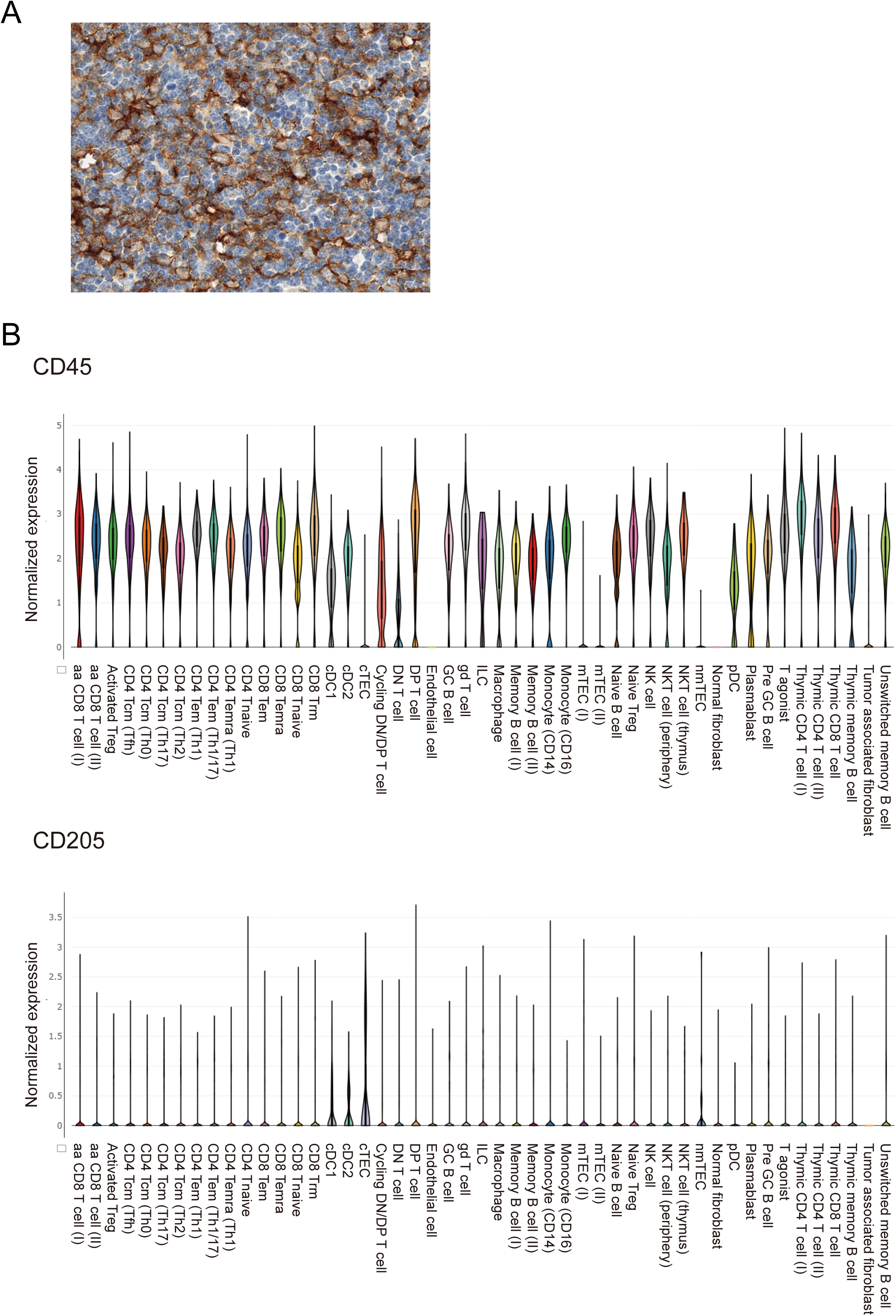
Expression of CD45 and CD205 in thymoma tissue. (A) Immunohistochemistry of CD205 in type B1 thymoma. (B) Droplet-based single-cell RNA sequencing (scRNA-seq) analysis of type B1 and B2 thymoma cells. Violin plots depicting expression levels of *CD45* and *CD205*.

**Supplemental Figure 2.**
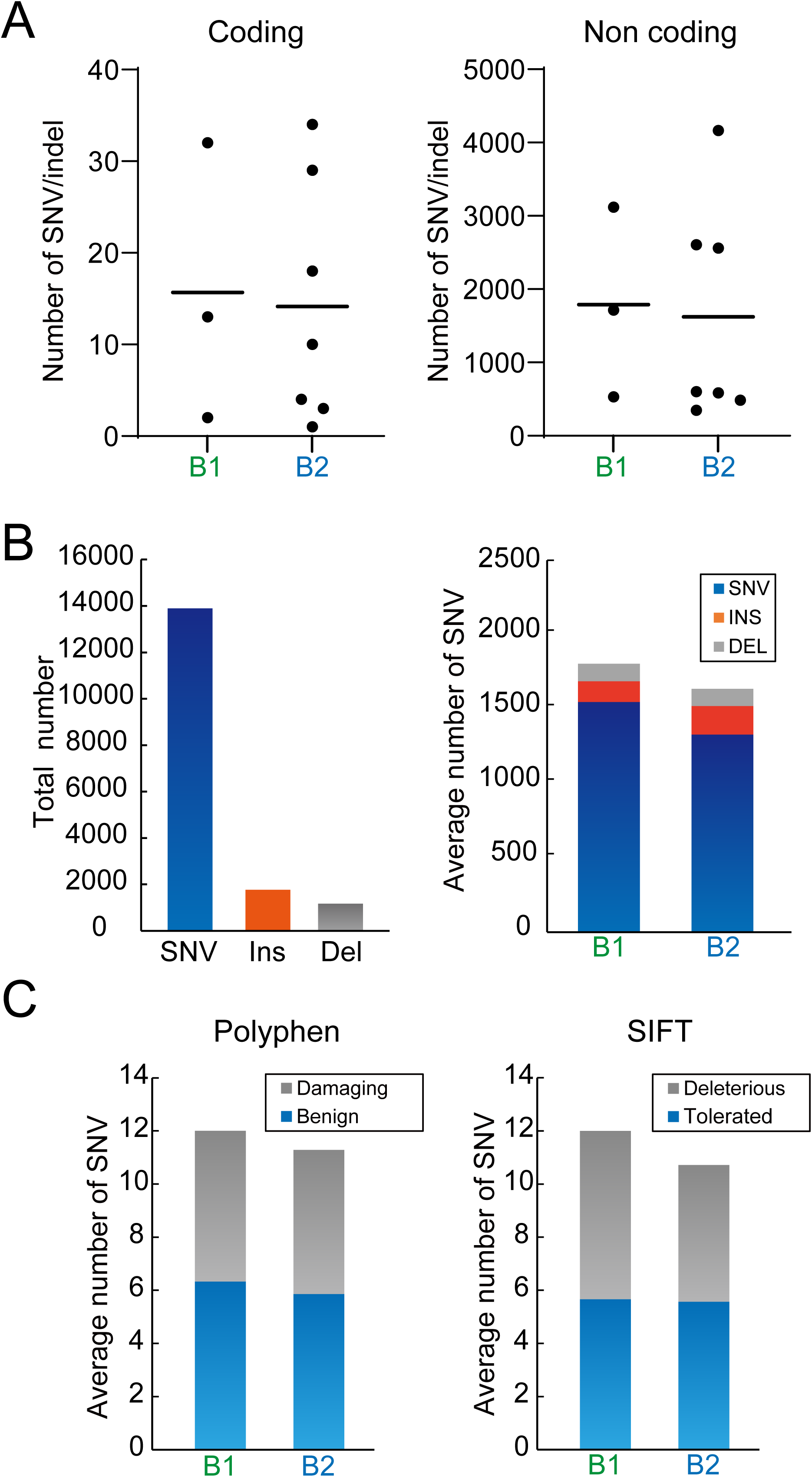
Comparison of somatic alterations between type B1 and B2 thymomas. (A) Coding and non-coding regions. (B) SNV, insertion, and deletion. (C) Polyphen and SIFT analysis.

**Supplemental Figure 3.**
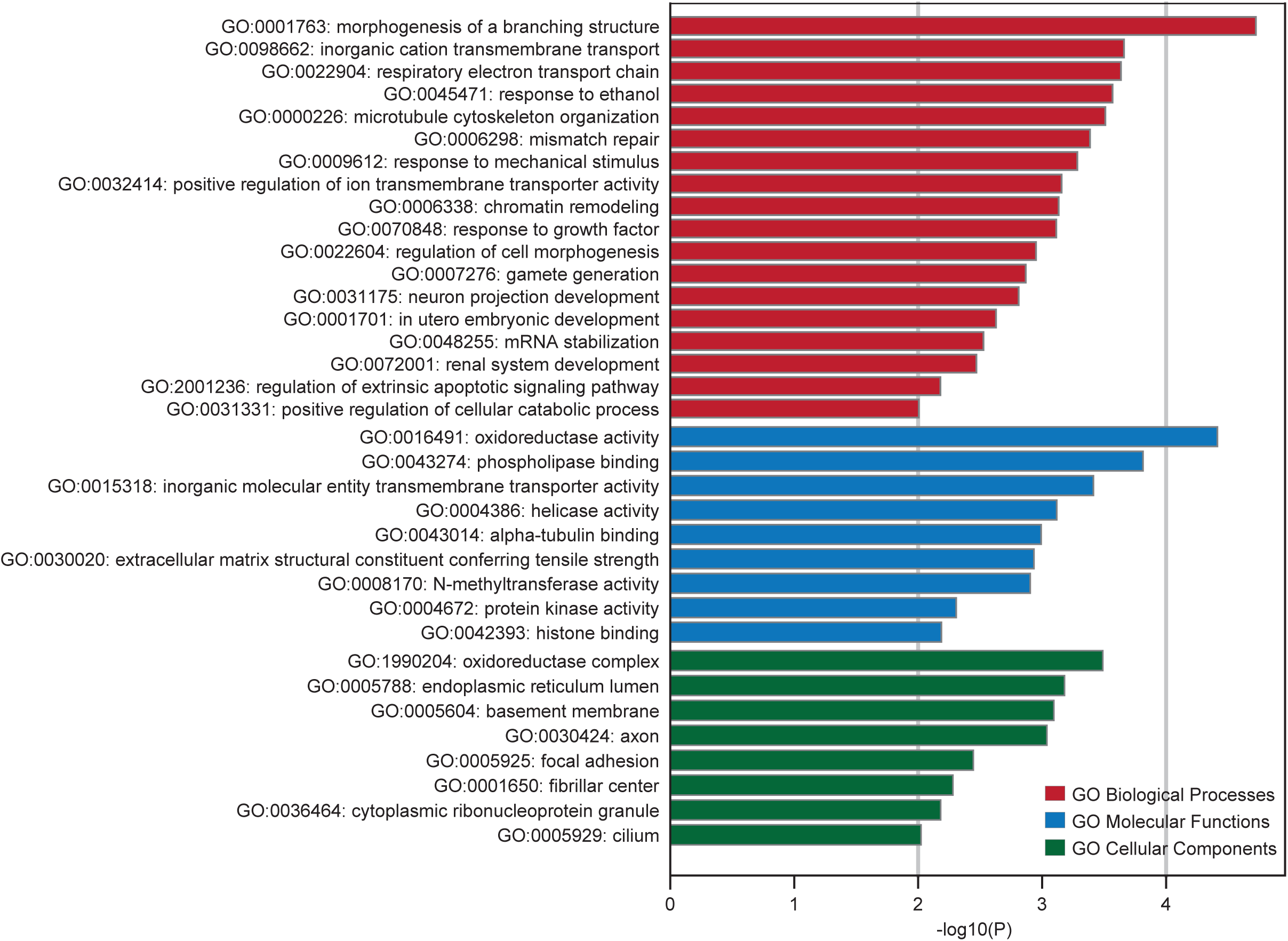
GO analysis. GO analysis for biological process, cellular component, and molecular function terms was performed on mutated genes; all terms with P < 0.01 are shown.

**Supplemental Figure 4.**
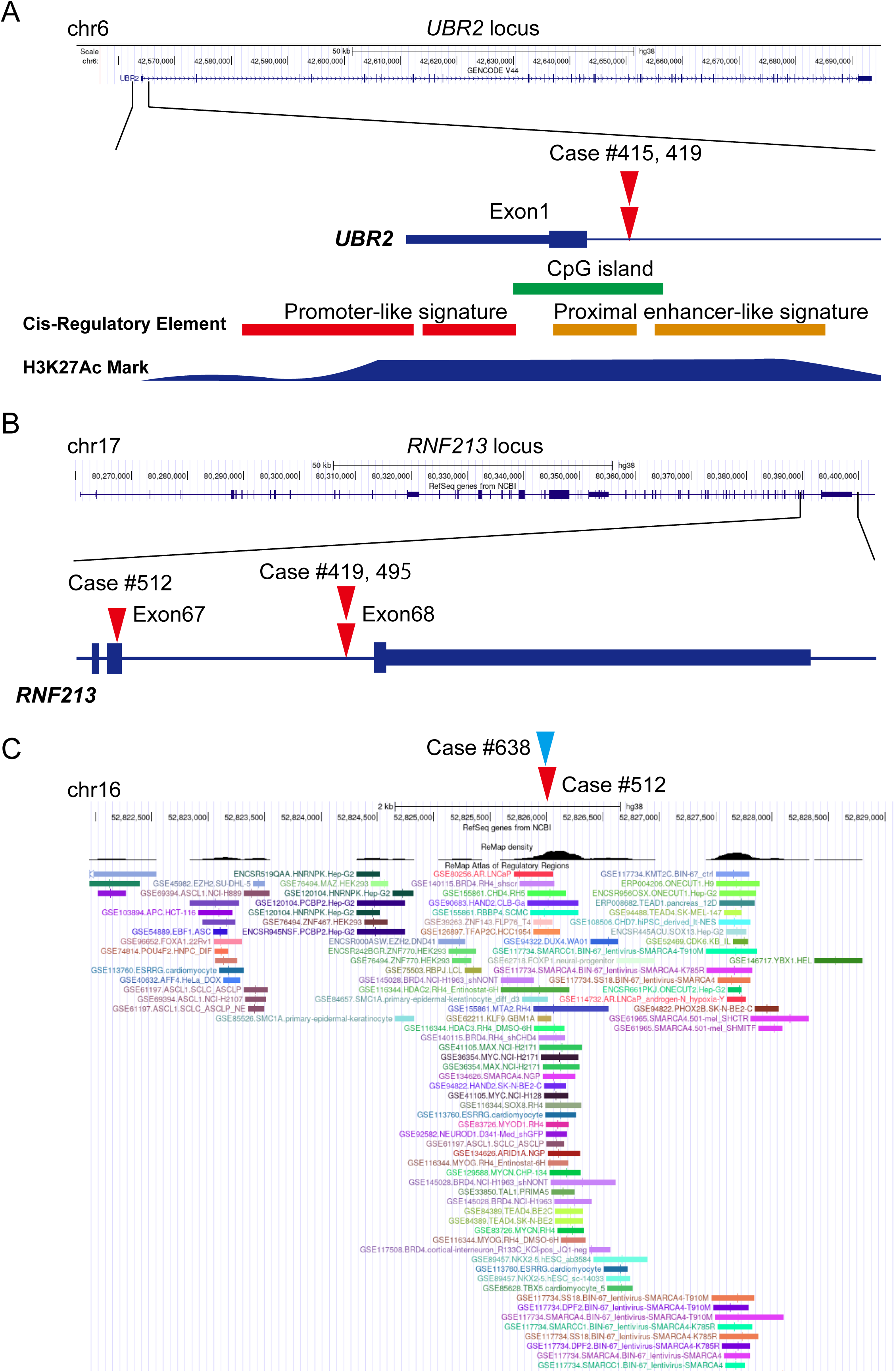
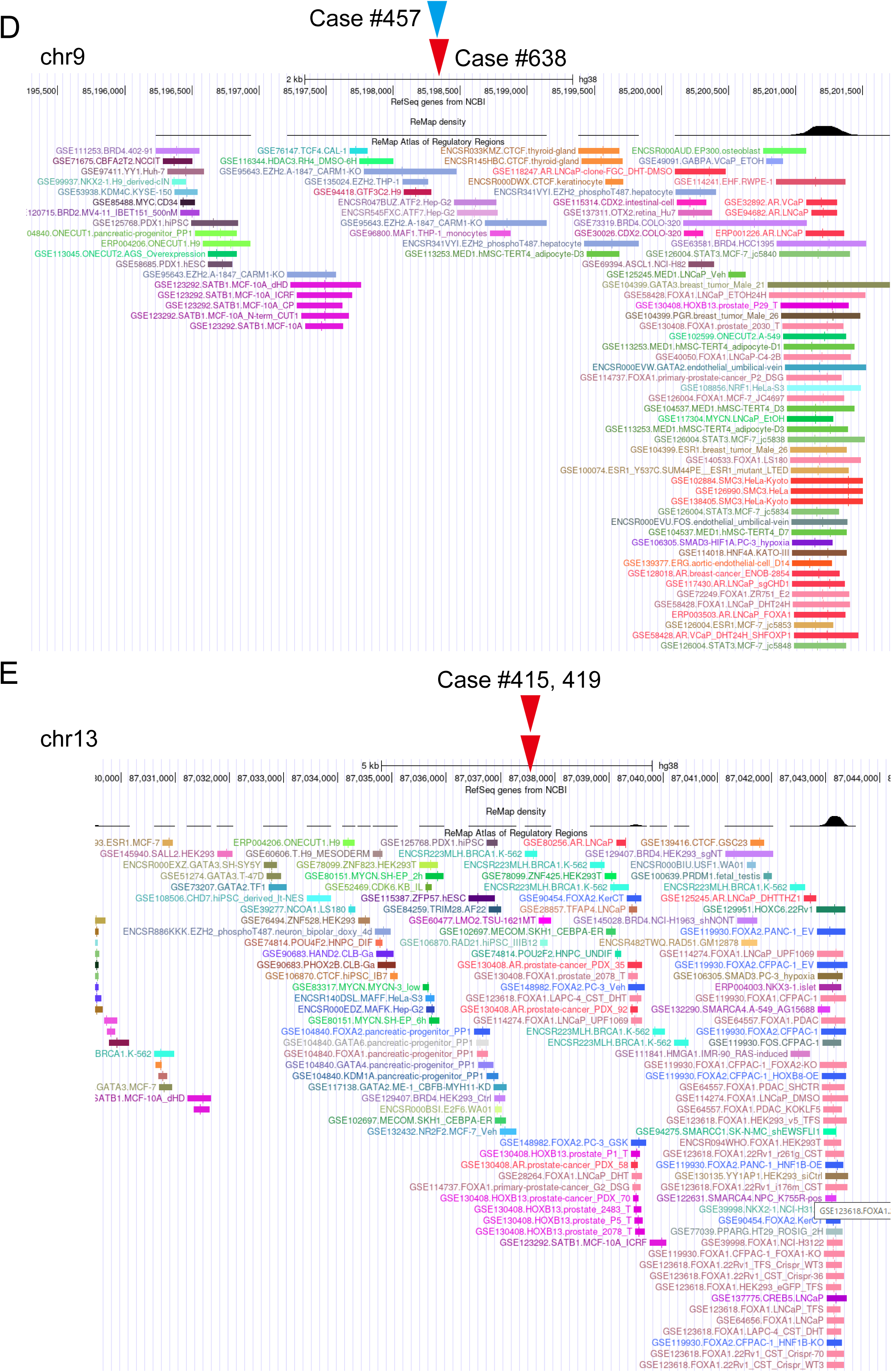
Recurrent non-coding mutations.

**Supplemental Figure 5.**
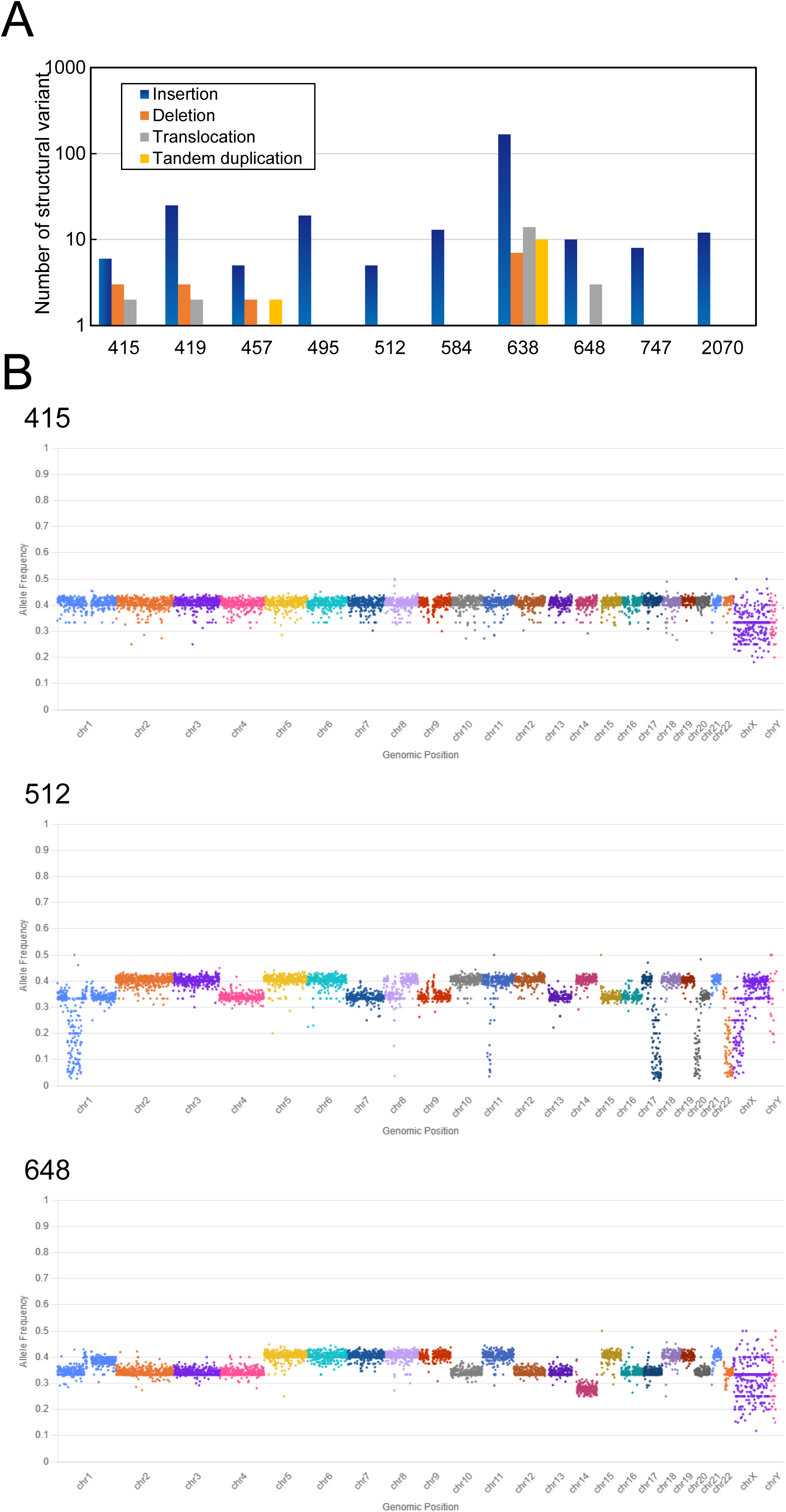
Copy number and structural variants in Type B1 and B2 thymoma. (A) Number of copy number and structural variants. (B) B-allele frequency for CNVs generated from DRAGEN somatic (v.4.2.7).

**Supplemental Table 1. Patient background.**

**Supplemental Table 2. Evaluation of deep whole-genome sequencing quality metrics.**

**Supplemental Table 3. The list of coding mutations.**

**Supplemental Table 4. The list of recurrent non-coding mutations.**

**Supplemental Table 5. The list of large copy number alterations.**

**Supplemental Table 6. The list of translocations.**

**Supplemental Table 7. The list of mitochondria DNA mutations.**

